# Vaxrank: A computational tool for designing personalized cancer vaccines

**DOI:** 10.1101/142919

**Authors:** Alexander Rubinsteyn, Isaac Hodes, Julia Kodysh, Jeffrey Hammerbacher

## Abstract

Therapeutic vaccines targeting mutant tumor antigens (“neoantigens”) are an increasingly popular form of personalized cancer immunotherapy. Vaxrank is a computational tool for selecting neoantigen vaccine peptides from tumor mutations, tumor RNA data, and patient HLA type. Vaxrank is freely available at www.github.com/openvax/vaxrank under the Apache 2.0 open source license and can also be installed from the Python Package Index.

## 1 Introduction

Mutated cancer proteins recognized by T-cells, known as “neoantigens”, are considered an essential component of a tumor-specific immune response (Finnigan *et al.*, 2015; Gubin *et al.*, 2015; Schumacher and Schreiber, 2015). Therapeutic vaccination against neoantigens is an emerging cancer therapy that attempts to mobilize an antigen-specific immune response against mutated tumor proteins (Türeci *et al.*, 2016; Zhang *et al.*, 2017). Since few tumor mutations are shared between patients, neoantigen vaccines must be personalized. A common approach for achieving personalization is high-throughput sequencing of tumor and normal patient samples followed by in-silico prioritization of mutated peptides that are likely to be presented on the surface of tumor cells by MHC (major histocompatibility complex) molecules.

Vaxrank is a tool for selecting mutated peptides for personalized therapeutic cancer vaccination. Vaxrank determines which peptides should be used in a vaccine from tumor-specific somatic mutations, tumor RNA sequencing data, and a patient’s HLA type. These peptides can then be synthesized and combined with an adjuvant to attempt to elicit an anti-tumor T-cell response in a patient.

The sequence of each mutated protein is determined by assembling variant RNA reads. Mutant protein sequences are ranked using a scoring system which seeks to satisfy two objectives: choosing mutations that are abundant in the tumor and choosing those whose translated amino acid sequences contain likely MHC ligands. Additionally, Vaxrank considers surrounding non-mutated residues in a peptide to prioritize vaccine peptide candidates and to improve the odds of successful synthesis.

Vaxrank was designed for and is currently being used in the Personalized Genomic Vaccine Phase I clinical trial at the Icahn School of Medicine at Mount Sinai (NCT02721043) (Rubinsteyn *et al.*, 2016a).

**Figure 1:**
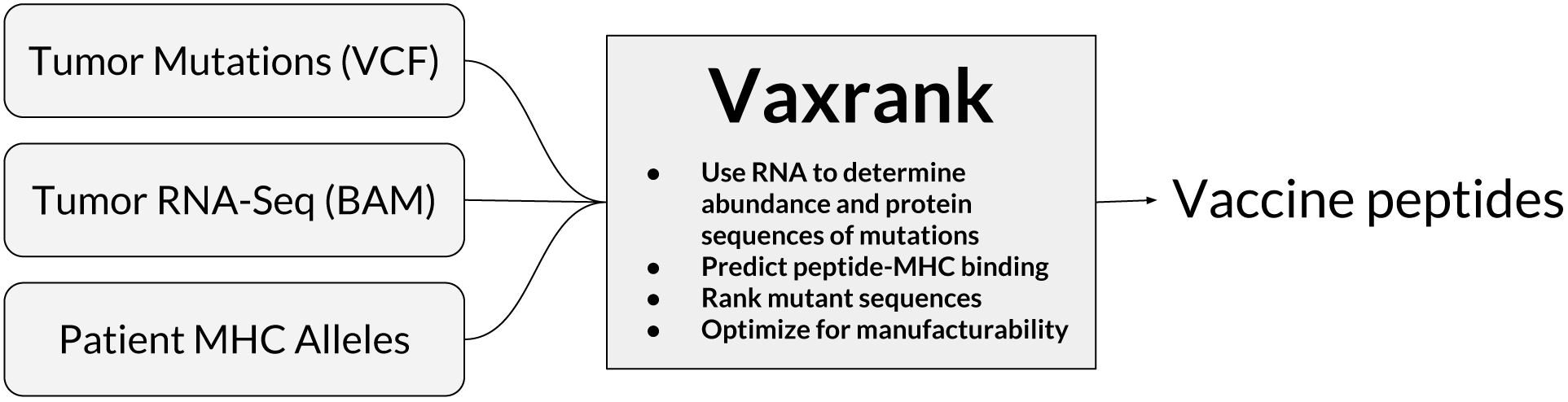
Users provide tumor mutations, tumor RNA sequence data, and patient HLA type. These are used to determine mutant protein sequences and rank them according to expression and predicted MHC affinity.

## 2 Running Vaxrank

To generate a Vaxrank vaccine report, the user must provide one or more files containing somatic variants (in VCF, MAF, or JSON format), aligned tumor RNA-seq reads (as an indexed BAM), and the HLA alleles to be used for MHC binding prediction:

`vaxrank`
`--vcf somatic-variants.vcf`
`--bam tumor-rna.bam`
`--mhc-predictor netmhc`
`--mhc-alleles H2-Kb,H2-Db`
`--mhc-peptide-lengths 8-10`
`--vaccine-peptide-length 21`
`--output-pdf-report vaccine-peptides.pdf`

Somatic variants can be specified using the `--vcf` option. Vaxrank also requires a BAM file of aligned tumor RNA reads `(--bam)`. This allows Vaxrank to both quantify tumor expression of each mutation and to phase adjacent variants when reconstructing a mutated coding sequence.

The `--mhc-predictor` argument controls which program is used to predict the affinity between a peptide-MHC pair. Vaxrank supports the use of locally installed instances of NetMHC (Andreatta and Nielsen, 2016), NetMHCpan (Nielsen *et al.*, 2007), NetMHCcons (Karosiene *et al.*, 2012), MHCflurry (Rubinsteyn *et al.*, 2016b), or a variety of web-based predictors through IEDB (Vita *et al.*, 2015).

Vaxrank’s output can be formatted as PDF, plain-text, HTML, or an Excel spreadsheet. The output lists variants in ranked order along with vaccine peptide(s) containing that variant, predicted MHC ligands, number of supporting RNA reads, and sequence properties that affect manufacturability. A larger list of options for input data, filtering, and output formats can be seen by running vaxrank `--help`. Documentation is available online at wwvaxrank.readthedocs.org

## 3 Ranking Mutations

A patient’s coding mutations are ranked according to a score that combines each mutation’s degree of expression and aggregate affinity of overlapping mutant peptides for that patient’s MHC alleles.

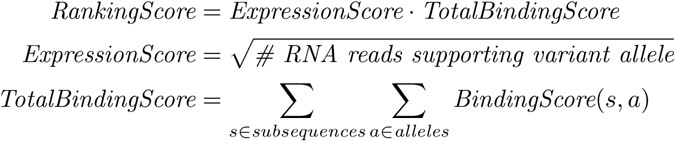

The *BindingScore* function is, by default, a logistic transformation of the peptide-MHC binding affinity that loosely approximates the probability of T-cell response (Sette *et al.*, 1994). Alternatively, binding predictions can be scored using an affinity threshold (commonly ≤500nM) or a threshold on the percentile rank of the affinity. Only subsequences which overlap mutant residues and do not occur in the reference proteome are considered as part of the *TotalBindingScore*.

It is important to note that there is at best a loose relationship between *RankingScore* and immunogenicity. Intracellular factors such as antigen processing are not captured by Vaxrank’s scoring logic. More importantly, even if a particular peptide is presented by a patient’s MHCs, it is not currently possible to predict whether it will generate a cytotoxic T-cell response. Further work is required to quantify the accuracy of Vaxrank’s ranking algorithm.

## 4 Manufacturability

Vaxrank was designed under the assumption that its output will be used to make long peptides, due to their favorable immunological properties (Rosalia *et al.*, 2013). Unfortunately, long peptides are also more difficult to synthesize using traditional solid phase chemistry (Bodanszky, 1988). To avoid known difficulties in synthesis, Vaxrank selects a window of amino acids around each mutation that minimizes the following undesirable properties:

1. total number of cysteine residues
2. *max*(0, mean hydrophobicity of 7 residues at C-terminus)
3. *max*(0, mean hydrophobicity of any 7 amino acid window)
4. glutamine, glutamic acid, or cysteine at N-terminus
5. cysteine at C-terminus
6. proline at C-terminus
7. asparagine at N-terminus
8. total number of asparagine-proline bonds

Manufacturability optimization does not affect the ranking of mutations but is only used for selecting which surrounding residues should be included. In cases where a mutation spans a “difficult” sequence (e.g. long hydrophobic stretch), minimizing these criteria may fail to salvage manufacturability.

## Supplemental Materials

### Installing Vaxrank

Vaxrank can be installed using pip:

`pip install vaxrank`

This will install the Vaxrank library, along with all of its dependencies, including PyEnsembl, Varcode, Isovar, and MHCtools.

To generate PDF reports, you first need to install wkhtmltopdf. On Mac OS X this can be done by running:

`brew install Caskroom/cask/wkhtmltopdf`

Vaxrank uses PyEnsembl for accessing information about the reference genome. You must install an Ensembl release corresponding to the reference genome associated with the mutations provided to Vaxrank.

The latest supported release for GRCh38 is Ensembl 87:

`pyensembl install --release 87 --species human`

The last release for GRCh37 was Ensembl 75:

`pyensembl install --release 75 --species human`

If using Vaxrank for a large number of mutations, it is recommended to locally install an MHC binding predictor such as NetMHCpan or MHCflurry. NetMHCpan can be downloaded from /www.cbs.dtu.dk/services/NetMHCpan/, whereas MHCflurry can be installed by running the following commands:

`pip install mhcflurry`

`mhcflurry-downloads fetch`

### Basic Usage

`vaxrank \`

`--vcf somatic-variants.vcf \`

`--bam tumor-rna.bam \`

`--mhc-predictor netmhc \`

`--mhc-alleles A*02:01,A*02:03 \`

`--mhc-epitope-lengths 8 \`

`--padding-around-mutation 5 \`

`--vaccine-peptide-length 25 \`

`--output-ascii-report vaccine-peptides-report.txt`

This tells Vaxrank to:

- load mutations from the input VCF file somatic-variants.vcf
- look for evidence of expression for mutations in the RNA BAM file tumor-rna.bam
- predict MHC binding of each possible 8mer peptide overlapping expressed mutations using the NetMHC prediction algorithm with the A*02:01 and A*02:03 MHC alleles
- choose protein vaccine candidates composed of 25 amino acids
- write top ranked variants with their associated vaccine proteins to `vaccine-peptides-report.txt`

You can read the complete Vaxrank documentation at http://vaxrank.readthedocs.io.

### RNA Options

Vaxrank uses the Isovar library to identify RNA reads supporting each genomic variant and assemble them into a mutant coding sequence. The assembly algorithm works by iteratively extending candidate sequences using overlapping reads. Determining the coding sequence from RNA reads, rather than simply substituting a variant into a reference transcript, Vaxrank is able to capture local phasing of variants. There are several Isovar options which are exposed as commandline flags in Vaxrank:

- `--min-mapping-quality` Reads which mapping quality below this value are ignored when gathering evidence of variant expression.
- `--use-duplicate-reads` Duplicate reads are normally ignored by Isovar/Vaxrank since they can inflate estimated abundance.
- `--min-alt-rna-reads` Controls the minimum number of RNA reads supporting a variant required to include that variant in the output report. Ignore variants with fewer than this number of supporting RNA reads.
- `--min-variant-sequence-coverage` Assembly of overlapping RNA reads may result in variable coverage across the assembled transcript fragment. This option allows the user to trim portions of the sequence that fall below some desired number of reads, creating a trade-off between assembled sequence and minimum coverage for each nucleotide.
- `--max-reference-transcript-mismatches` The reference transcriptome is used to determine the reading frame for an assembled sequence. Any transcript with more than this number of mismatches is discarded when trying to determine the reading frame.
- `--include-mismatches-after-variant` Make mismatches after the variant locus count toward the `--max-reference-transcript-mismatches` filter. This is normally disabled since technically only the sequence leading up to a variant should affect its reading frame.–
- `--min-transcript-prefix-length` Number of nucleotides before the variant we try to match against a reference transcript. Values greater than zero exclude variants near the start codon of transcripts without 5’ UTRs.

### Performance

The MHC binding prediction algorithm used by Vaxrank can profoundly influence how long it takes to determine results for a patient. To give a rough sense of Vaxrank’s performance, we generated the following timing results on a 2013 Macbook Air. The inputs were patient data with 521 somatic variants, 83.7M aligned tumor RNA reads, and peptide predictions for lengths 8-11.

| Predictor | Time |
| --- | --- |
| NetMHC 4.0 | 42 seconds |
| NetMHCpan 3.0 | 2 minutes, 17 seconds |
| MHCflurry 0.9.1 | 44 seconds |

